# Increased frequencies of IgH locus suicide recombination points on Chronic Lymphocytic Leukemia with low rate of AID related somatic mutations, cMYC overexpression and short telomeres

**DOI:** 10.1101/2022.03.06.482661

**Authors:** Israa Al Jamal, Milène Parquet, Hend Boutouil, Kenza Guiyedi, David Rizzo, Marine Dupont, Mélanie Boulin, Chahrazed El Hamel, Justine Lerat, Saïd Aoufouchi, Samar al Hamaoui, Nehman Makdissy, Jean Feuillard, Nathalie Gachard, Sophie Peron

## Abstract

In normal activated B-cells, Activation Induced-cytidine deaminase (AID) is absolutely required for immunoglobulin (Ig) class switch recombination (CSR) and IGHV somatic hypermutation (SHM). AID is also implicated in the Locus Suicide Recombination (LSR) of the Ig heavy (IgH) locus, resulting in the deletion of the *IgH* constant part. Chronic lymphocytic leukemia (CLL) is an indolent non-Hodgkin B-cell lymphoma characterized by tumor CLL B-cells that weakly express a B cell receptor (BCR) on their surface. The great majority of CLL tumor B-cells are non class-switched. Searching for abnormalities of *IgH* locus recombination in CLL, we investigated CSR and LSR in samples from CLL patients (N=47) with high blood tumor cell infiltration (>98%) and in healthy volunteers (HV) as controls (N=9). LSR was detectable at comparable levels in both HV and CLL groups. CSR counts were decreased in CLL samples as expected. As distribution of LSR counts was bimodal, we separated CLL patients in two groups so called LSR^High^ and LSR^Low^. LSR^High^ CLLs exhibited very weak AID expression and low mutation rate of IgHV region and of the AID off-target PIM1 gene. LSR junction diversity, evaluated using the Shannon index, was increased in LSR^High^ CLLs suggesting that LSR was on-going in these cells. Also, shorter telomeres were observed in LSR^High^ CLLs suggesting an increased number of past mitosis. Consistently, increased levels of cMYC expression were detected in LSR^High^ CLLs and treatment free survival of these cases was markedly decreased. We hypothesized that LSR in LSR^High^ CLLs is AID independent and could be due to DNA lesion and inaccurate DSB repair within the IgH locus, which could be accessible to recombination machinery due to increased *IgH* locus transcription. Altogether, our results indicate the accessibility of *IgH* locus and the proliferation increase LSR rate in LSR^High^ CLLs could be related to cMYC resulting in shorter treatment free survival of patients and point on an AID independent mechanism of IgH recombination.

## Introduction

In normal B-cells, Activation Induced-cytidine deaminase (AID) is the key enzyme for class switch recombination (CSR) and IGHV somatic hypermutation (SHM) in activated B-cells ^6,7^. AID is also implicated in another IgH rearrangement, the Locus Suicide Recombination (LSR). LSR occurs in activated B-cells and recombines the IgH locus between the switch μ (Sμ) region and one 3’a2 regulatory region (3’α2RR) of the *IgH* locus. The 3’a2RR contains enhancers (HS3, HS1.2, and HS4) and controls *IgH* locus transcription necessary for IgH expression^8^. LSR results in the complete deletion of the cluster of IgH constant genes. When LSR hits the active *IgH* locus, it induces the loss of BCR expression and the death of the B-cell^9–11^.

Being still incurable, chronic lymphocytic leukemia (CLL) is an indolent non-Hodgkin B-cell lymphoma of the elderly that results from the expansion of malignant CD5+ B cells. CLL B-cells weakly express the B cell receptor (BCR) on cell surface, that BCR composed, in the vast majority of cases, of immunoglobulins (Ig) of the mu (μ) and delta (δ) isotypes (IgM^+^IgD^+^) ^12^. Ig class-switched CLLs are rare^13^. In IgM^+^IgD^+^ CLLs, when present, genuine switched clonal cells are unfrequent ^14–16^. Ig class-switched CLL cells were shown to express AID at higher levels^15,17–19^. In CLL, AID-dependent CSR is in fact dissociated from SHM. Indeed, AID expression is increased in class-switched CLLs, which are predominantly unmutated (U-CLL) ^17^.More broadly, increased AID expression in CLL is not only associated with unmutated IGHV genes but also with unfavorable cytogenetic aberrations and poor prognosis^20^. This raises the question of abnormalities in the Ig gene recombination machinery in this B-cell cancer.

Here, we investigated whether CLL cells are induced to LSR like they can undergo to CSR. Our results led us to point on LSR^High^ CLLs which are biologically different and suggest an AID independent mechanism of *IgH* locus recombination.

## Methods

### Human materials and ethics

The project was conducted according to the guidelines of the Declaration of Helsinki. CLL Peripheral Blood Mononuclear Cells PBMC are from CRBioLim from Limoges Hospital, CHU Dupuytren (authorization: DC-2008-604, AC-2016-2758, and AC-2019-3418). Tonsils were obtained from children scheduled for elective tonsillectomy and were obtained from CRBioLim (authorization: DC-2008-604, AC-2018-3157). Normal PBMC from healthy volunteers were collected through research project approved by CPP Sud Méditerranée I (N°. 2021-A00778-33).

### DNA extraction

PBMC DNA were extracted using phenol/chloroform method. CLL DNA were extracted using kit QiAmp DNA Blood (51104 Qiagen).

### Count of Human CSR and LSR junctions

CSR and LSR junctions were amplified as previously described ^11^ and used to prepare next generation sequencing (NGS) libraries (Ion Xpress™ Plus Fragment Library Kit, life technologies, thermofisher, 447269) sequenced with ion proton or S5 chip (life technologies Results are collects as FastQ format then proceeds to the bioinformatics using CSReport ^21^. Shannon diversity index (*H*) was used to estimate the diversity of the samples and was calculated considering the number of reads (*ni*) for each particular LSR junction and the total read number (*N*) of LSR junction. 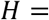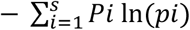 with 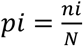.The *H* value varies from 0 when LSR junction diversity shrinks due to clonal dominance to high values for samples with higher diversity.

### IgHV sequence analysis

Amplification of V, D, and J rearranged genes was performed using the Biomed-2 strategy with FR1 and FR2 primers and sequence analysis were performed as previously described ^22^.

### Flow cytometry analysis

Immunophenotyping was done on navios-Beckman coulter (B83535) using: CD5-APC (Beckman coulter PN a60790, clone BL1a), CD19-ECD (Beckman coulter A07770, clone J3-119) and dual-color reagent Anti-human Kappa light chain/Anti-Human Lambda light chains/RPE (Dako, FR481 X0935). Results were analyzed with Kaluza software version 2.1 (Beckman coulter).

### RNA extraction, cDNA synthesis and quantitative real time PCR

Total RNA was isolated using TRIZOL reagent (TRIzol™ Reagent, 15596018) and reverse transcripted (Advantage RT-for-PCR kit Applied Biosystems™, thermofisher 4368814/10400745). qRT-PCR were performed with the SYBR Green PCR mix (SensiFast hi ROX syber green BIO820025) for different interest targets (supplemental table 3) or with Taqman PCR mix (SensiFast Probe Hi-Rox kit BIO820025) and cMYC probe (4331182 Hs00905030_m1, thermofisher). Normal centroblasts were sorted as described ^23^ from tonsils.

### Relative Telomere length assay (RTL)

25ng of DNA of CLL patients and normal PBMC used in triplicate to perform the analysis of relative telomere length by qPCR as already described ^24^.

### Mutation analysis of PIM1

We amplified the PIM1 exon4 segment DNA with specific primers (Supplemental table 3) as this exon contains a base pair shown to be the AID-targeted in mouse model of CLL and in human CLL ^25^. Amplification used Phusion High fidelity Taq polymerase (Thermo Scientific, F-530XL) and were used to build NGS libraries and submitted to NGS as describes above. For each library analysis was done by alignment of sequenced reads with the reference sequence NM_002648.4 using the Torrent Mapping Alignment Program (TMAP) for Ion Torrent Data and Super-maximal Exact Matching algorithm ^26^. The resulting BAM files were processed to generate per-base nucleotide count tables files consisted of matrices with *n* lines × 4 columns (*n* is the length of the sequenced DNA) and the columns correspond to nucleotides (A, C, G, and T). The consensus sequence is the most frequently read nucleotides and corresponds to the sequence reference. Count of mutated bases was calculated by addition of numbers of sequenced bases different from nucleotide that is sequenced the more frequently.

### Statistical analysis

Graphs, histograms, curves, and statistical analysis are designed using graph pad 6-prism software.

## Results

Searching for abnormalities of *IgH* locus recombination, we have analyzed DNA samples from CLL patients (N=47) with more than 98% blood tumor cell infiltration (Sup figure 1). Controls consisted in healthy volunteers (HV) (N=5 for CSR and N=9 for LSR). As shown in figure 1A and 1B, both CSR and LSR were detectable. As expected in this IgM+ IgD+ B-cell cancer, CSR levels were much lower in CLL than in HV samples and none of CLLs exhibited increased CSR counts (Figure 1A). Even if at low levels, LSR was found at comparable levels in both HV and CLL (Figure 1B). LSR was undetectable in only 3/47 (6.3%) CLL patients. Oppositely, some patients had increased LSR counts when compared to HVs (Figure 1B). Indeed, distribution of LSR counts was bimodal with a valley at 27. That value being also the mean of LSR counts in HVs (Sup figure 2), we separated CLL patients in two groups so called LSR^High^ (12/47 patients = 26%), and LSR^Low^ (35/47 patients = 74%) (Sup table 1). Noteworthy, CSR counts were not significantly different in both groups (Figure 1C). According to the same methodology, with a threshold of 800 CSR counts, we also separated CLL patients in CSR^Low^ and CSR^High^. As shown in supplementary table 2, most of LSR^Low^ CLLs were also CSR^Low^. In contrast, even if numbers are small, the LSR^High^ status does not seem to be related to the CSR^High^ status. Indeed, the positive predictive power (ppp) of LSR^High^ to predict CSR^High^ is 5/12 (42%) while the ppp of CSR^High^ to predict LSR^High^ is 7/15 (47%). Consistently, correlation between CSR count and LSR count was poor (correlation coefficient r=0.2, not shown). This indicates that increased Locus Suicide Recombination was poorly related to class-switch in CLL.

**Figure 1:**
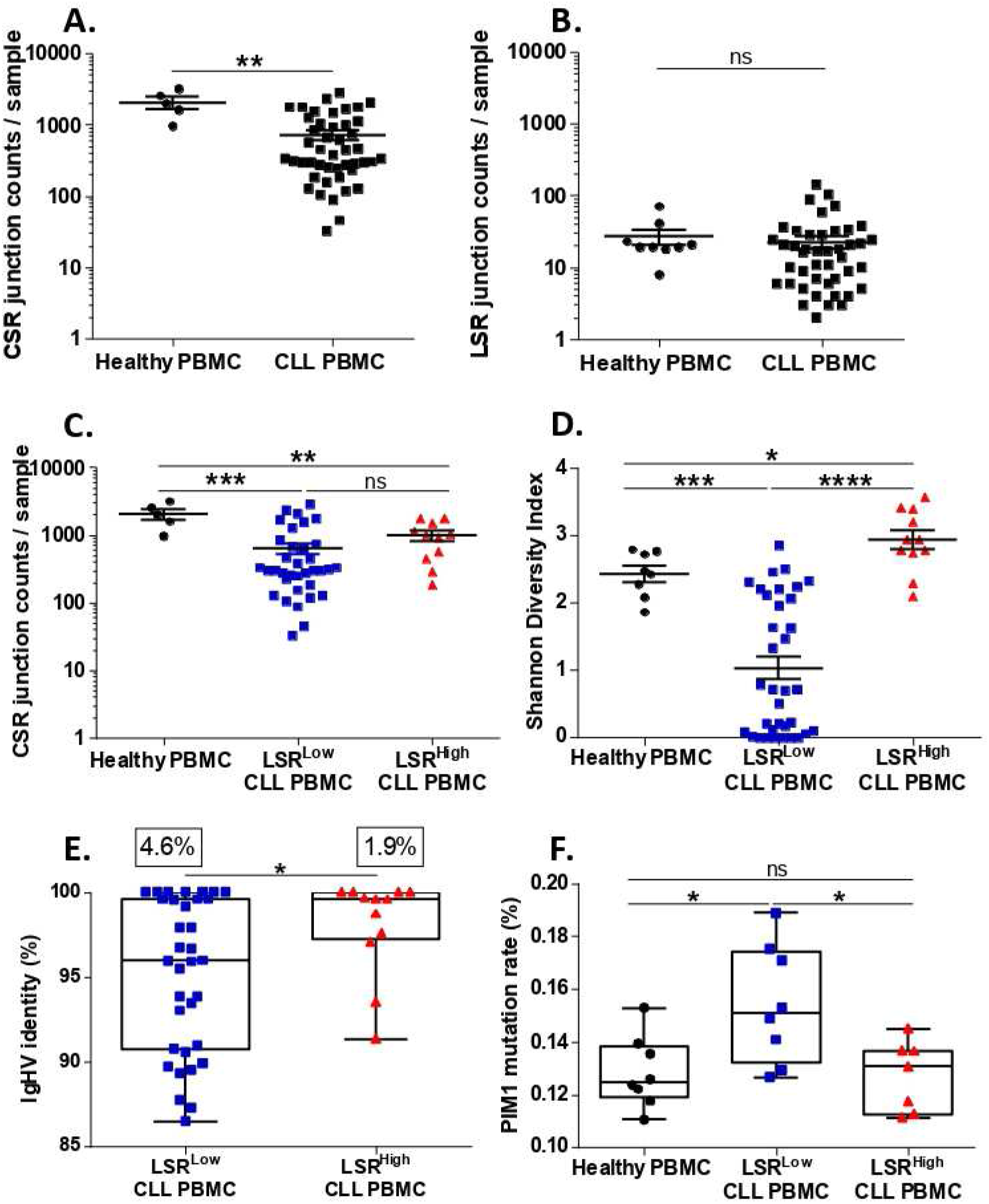
Locus Suicide Recombination (LSR) is detectable in CLL and points on patients with increased LSR rate. **A.** CSR junction counts analyzed by NGS and CSReport in CLL patients (N=47, 32986 junctions) is lower than in HV PBMC (N=5, 10301 CSR junctions). **B**: LSR junctions were at comparable levels in both CLL samples (N=47, 1060 LSR junctions) and HV PBMC (N=9, 239 LSR junctions). **C.** CSR counts were not significantly different in both LSR^Low^ (N=35, 22247 junctions) and LSR^High^ (N=11, 10739 junctions). **D.** Shannon index indicates higher LSR junction diversity in LSR^High^ CLL samples (N=11) than in LSR^Low^ (N=35) CLLs and healthy PBMC (N=9). **E.** Graph represents the percentage of identity in IgHV segment, in LSR^Low^ (N=34) low % of identity detected compared to the high homology of IgHV segment in LSR^High^ (N=12) groups. The mean frequency of Somatic Hyper Mutation (SHM) in the two groups of CLL is represented in the box. **F.** NGS of PIM1, AID off-target gene, and mutation frequency analysis show high level of mutation frequency in LSR^Low^ (N=8) compared to healthy PBMC (N=8) and LSR^High^ (N=7). Graphs represented mean ± SEM. Statistical analysis were performed using Unpaired T test ns: non-significant, *P<0, 05, **P<0.01, ***P<0.001 and ****P<0.0001.

**Figure 2:**
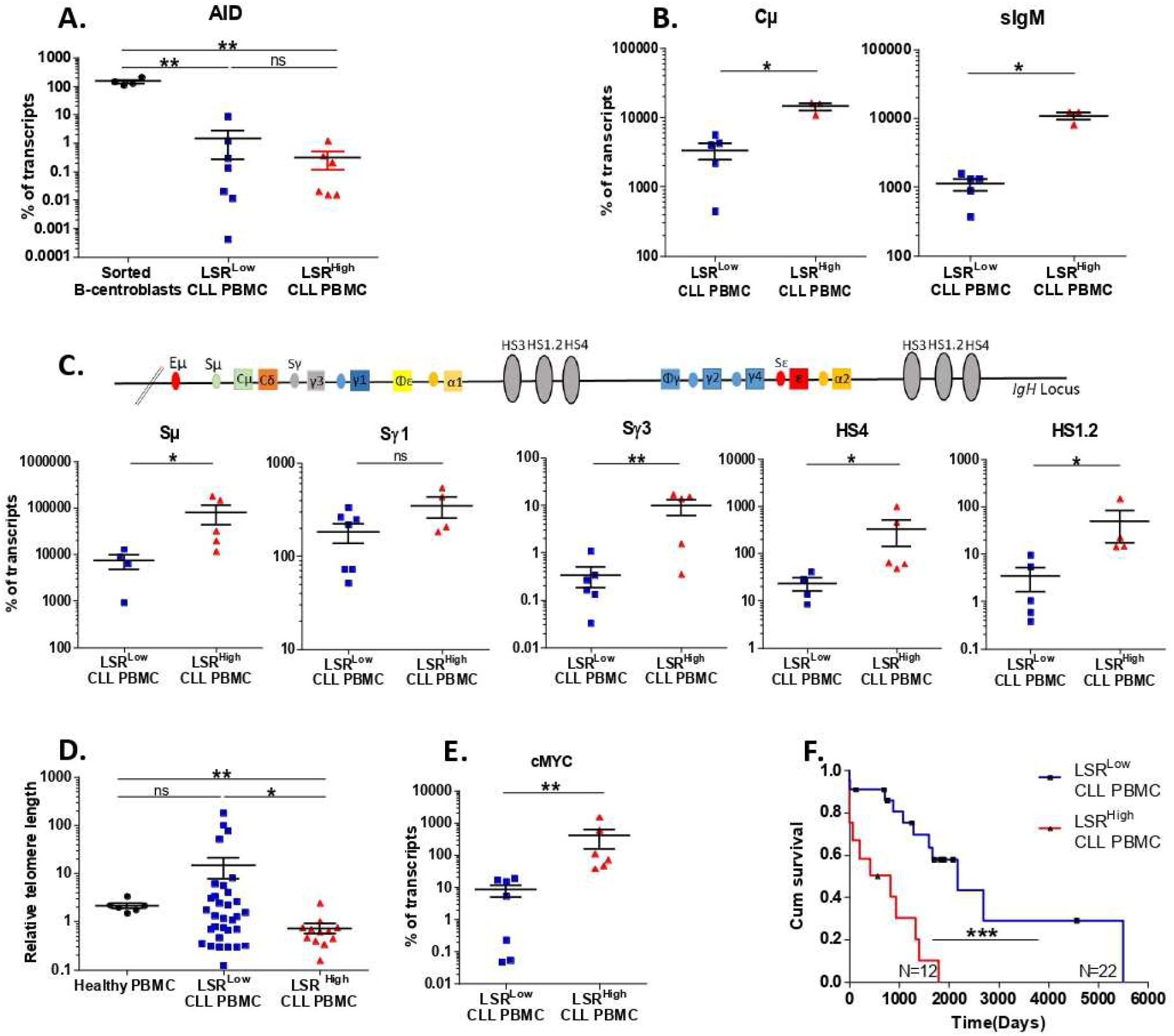
LSR in CLL seems not AID-related in LSR^High^ patients. **A.** Quantification of AID transcripts, relatively to CD19 transcripts, are low in LSR^Low^ CLLs (N=7) and LSR^High^ CLLs (N=6) compared to normal B-centroblasts (N=4) used as positive controls. Quantification of productive transcripts (Cμ and Surface IgM (sIgM)) of *IgH* locus represented in (**B.)** and non-productive transcripts of *IgH* locus (Sμ, Sγ1, Sγ3, HS1.2 and HS4) represented in graphs (**C.**) in CLL LSR^Low^ (N=4 to 7) and LSR^High^ (N=3 to 5) CLLs. LSR^High^ exhibits high levels of IgH locus transcription in both productive and non-productive transcripts. **D.** Relative telomere length (RTL) was measured by specific qRT-PCR relatively to human beta globin gene (LSR^Low^, N=33; LSR^High^, N=12; healthy PBMC, N=6). Telomere length was significantly shorter in LSR^High^ comparet to LSR^Low^ and healthy PBMC. **E.** cMYC expression was higher in LSR^High^ (N=6) compared to CLL LSR^Low^ (N=7). **F.** Cumulative survival time (days) without treatment (TFS) for patients indicated shorter TFS in LSR^High^ CLLs. Graphs represented mean ± SEM. Statistical analysis was performed using Unpaired T test (A., B., C., D., E.) or Chi2 test (F.) ns: non-significant, *P<0, 05, **P<0.01, and ***P<0.001.

To further investigate LSR^High^ CLLs, we analyzed the diversity of the LSR junctions in both groups and in HVs using Shannon Index (Figure 1D). This index was significantly higher in LSR^High^ CLL samples than in LSR^Low^ CLLs and HVs, indicating an increased junction diversity. However, IgHV mutation rate was quite different in both groups. Indeed, LSR^High^ samples exhibited a stronger homology to IgHV reference sequences (Figure 1E). With the a threshold of 95% homology 10/12 (83%) LSR^High^ CLLs were none or weakly mutated while 20/35 (57%) LSR^Low^ CLLs were highly mutated (Fischer test, p=0.02). Consistently, we found that LSR^High^ patients exhibited low rate of AID off-target PIM1 mutations (Figure 1F, p=0.01). When compared to centroblasts sorted from benign reactive tonsils, AID expression was very low and at comparable levels in both LSR^Low^ and LSR^High^ CLL (Figure 2A).

Because diversity of LSR junctions in LSR^High^ CLLs is evocative of an on-going process, we analyzed the expression of productive and non-productive transcripts of the constant part of *IgH* locus, which we found at higher level in LSR^High^ CLLs compared to LSR^Low^ CLLs (Figure 2B and 2C), meaning that IgH locus DNA was accessible to the recombination machinery in these patients. As *IgH* locus recombination physiologically occurs in proliferating cells ^27^, we measured the relative telomere length. While being homogeneous in HVs, with a mean of 2.14, telomere lengths were very heterogeneous in LSR^Low^ patients, 13 of them (40%) with long or very long telomeres that should correspond to a very small number of past proliferation cycles. In contrast, all but one LSR^High^ patients homogeneously exhibited telomeres shorter than HVs that would reflect an increased proliferation rate (Figure 2D). Moreover, shorter telomeres in LSR^High^ CLLs was associated with increased expression of cMYC (Figure 2E). Finally, the Kaplan Meyer curves of Treatment Free Survival (TFS) show TFS of LSR^High^ CLLs was significantly shorter than the one of LSR^Low^ CLLs (≈14 months compared to ≈71 months; P<0.001) (Figure 2F).

## Discussion

In this study, we show that Locus Suicide Recombination (LSR) is detectable in most patients with CLL. We also point on patients in which LSR rate is increased.

In normal B-cells, both CSR and LSR depend on AID. Here, AID expression was indifferently low in both LSR^Low^ Low and LSR^High^ CLLs. Moreover, we did not find significant correlation between CSR and LSR counts suggesting that both processes were not related. LSR^High^ patients exhibited low mutation region of IgHV and PIM1 genes, an additional indication that LSR was not related to AID. Despite the absence of relationship between LSR and genetic markers of AID activity, increased levels of IgH transcription reflects the *IgH* locus accessibility to recombination events. Because LSR junction diversity was increased in LSR^High^ CLLs which evokes an on-going process, and IgH recombinations occur preferentially in proliferating cells, we studied the telomere length and show the LSR^High^ CLLs have constantly short telomeres, which suggests an increased number of past mitosis. Consistently, cMYC expression levels were increased in LSR^High^ CLLs. In agreement with the fact that decreased telomere length is a poor prognosis factor associated with genomic instability and TP53 mutations ^28^ we found that TFS of LSR^High^ patients was shorter. LSR could be induced in CLL proliferation centers in secondary lymphoid organs ^1^. It has been shown that some residual CSR can occur in absence of AID ^2^. This seems to result from random DNA breaks and inaccurate DSB repair as also implicated in chromosome translocations. We hypothesized that LSR in CLL could be due to similar inaccurate DSB repair related to increased proliferation rate and genetic instability. Indeed, CLL is known to harbor DNA repair alterations and to accumulate of DSB across the genome ^3–5^. This AID independent IgH locus recombination would explain why IgHV and PIM1 mutation rate was low in LSR^High^ CLLs.

Altogether, our results strongly indicate the accessibility of *IgH* locus and the proliferation in CLL patients with high rate and increased diversity of LSR junctions could be increased in cMYC dependent manner resulting in shorter survival and pointing on an AID independent mechanism of IgH recombination.

## Acknowledgements

This work was supported by grants from la Ligue Contre le Cancer (Comité de la Haute-Vienne, CD87). We are grateful to Dr. Villéger at the CRBioLim and to Dr. Vallejo and the INSERM CIC 1435, Dupuytren Hospital, Limoges, France, for providing human samples. IAJ is supported by Fondation pour la Recherche Medicale (FRM) and Association Libanaise (AZM and Saade, LASeR). KG is supported by Région Nouvelle Aquitaine and Délégation INSERM Nouvelle Aquitaine. MP is supported by Région Nouvelle Aquitaine and Université de Limoges. SP is a National Institute of Health and Medical Research (INSERM) investigator.

## Authorship Contributions

IAL performed investigation and methodology and participated in writing of the original draft. H.B, MP, KG, MB and MD participated in investigation and methodology.

DR, MD and NG participated in data curation.

CEH and JL allowed tonsils from patients undergoing tonsillectomy performed in Limoges Dupuytren hospital.

SA, SAH and NM participated in writing of the original draft.

JF, NG and SP led the conceptualization, data curation, funding acquisition, writing manuscript.

