## supplemental data for "Increased frequencies of IgH locus suicide recombination points on Chronic Lymphocytic Leukemia with low rate of AID related somatic mutations, cMYC overexpression and short telomeres"

**Supplemental table 1.** Numbers of Healthy volunteers and CLL patients tested and LSR junction counts and intervals (minimum-maximum) obtained in each group. Based on the mean of junction count obtained in healthy PBMC we divided the CLL cohort into two groups the first: $LSR^{Low}$ ≤27 junctions and $LSR^{High}$ > 27 junctions per sample.

| Count of LSR using CSReport | Healthy PBMC | $LSR^{Low}$ CLL PBMC | $LSR^{High}$ CLL PBMC |
|---|---|---|---|
| LSR positive samples / total samples (%) | 9/9 (100) | 32/35 (91) | 12/12 (100) |
| Mean of LSR junction (min-max) | 26.6 (8-71) | 10.2 (0-24) | 58.5 (29-145) |

**Supplemental table 2.** Repartition of patients between to two groups of LSR ($LSR^{Low}$, $LSR^{High}$) and CSR ($CSR^{Low}$, $CSR^{High}$). Statistical analysis was performed using Chi2 test *P<0.05.

| Samples * (P=0,05) Chi2 test | $LSR^{Low}$ CLL PBMC | $LSR^{High}$ CLL PBMC |
|---|---|---|
| $CSR^{Low}$ CLL PBMC | 27 | 5 |
| $CSR^{High}$ CLL PBMC | 8 | 7 |

**Supplemental Table 3.** Primers used in this study.

| Target and segment | | Primer name | Sequence 5'-3' |
|---|---|---|---|
| *IgH* locus Transcription | Sμ | Sμg | F:GGTGTGGGTTTTCACAGCTT |
| | | | R:CCTCACCAAGTCCACCAAGT |
| | Sγ1 | Sγ1-3 | F:CTGGGATGGAGAAGGGAAGG |
| | | | R:CTGGTCTCAAGCACACGTTC |
| | HS | HS1,2 | F:GAGTTTTCGGCATCTCTGGG |
| | | | R:ACAGATCAGAGCCCTCACAC |
| | | HS4 | F:GTGTGTCTGAGGGTGAGTGA |
| | | | R:ACACTGTCACACACTCCACA |
| | Sγ3 | Sγ3 | F:AGCTGTGCAACTGGAGTCCT |
| | | | R:TGAGCCACCTAATCCAAACC |
| | IgM total (Cμ) | hIgM CH1 | F :CAGAATGCGTCCTCCATGTG |
| | | | R :GGTGGACTTGGTGAGGAAGA |
| | Surface IgM | hsIgM-CH4-For | F: AAGAGGAATGGAACACGGGG |
| | | hsIgM-MB-Rev | R :ACAAGGTGACGGTGGTACTG |
| Telomere length | Telomere | Telomere A | CGGTTTGTTTGGGTTTGGGTTTGGGT TTGGGTTTGGGTT |
| | | Telomere B | GGCTTGCCTTACCCTTACCCTTACCCT TACCCTTACCCT |
| AICD transcription | AID | | F :GAGGCAAGAAGACACTCTGG |
| | | | R :GTGACATTCCTGGAAGTTGC |
| PIM1 mutation | PIM1 | | F :ATGAGTGGGTGGGGTGAGG |
| | | | R :ATCGAGCCAGGCGGCCC |
| Internal control | CD19 | | F:AGACTCCTTCTCCAACGGTA |
| | | | R:GGTCAGCTCTTCATCCTCGT |
| | Human Beta globin (hbg) | Hbg1 | GCTTCTGACACAACTGTGTTCACTAGC |
| | | Hbg2 | CACCAACTTCATCCACGTTCACC |

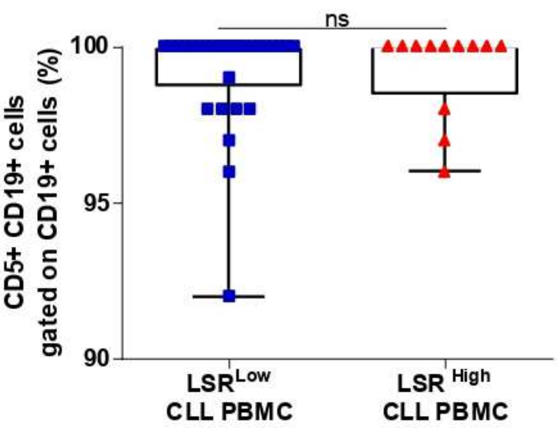


**Supplemental figure 1.** Blood tumor infiltration in the two CLL groups based indicated by the percentage of CD5+ CD19+ B cells gated on total CD19+ B cells.

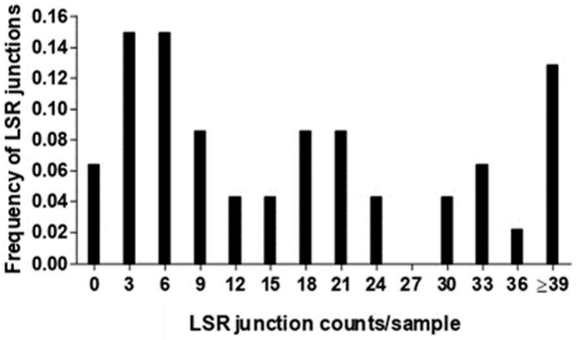


**Supplemental figure 2.** Distribution of LSR junction counts in CLL patients is bimodal with a valley at 27.

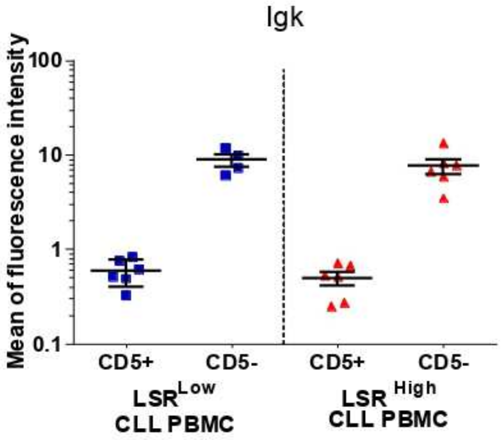


**Supplemental figure 3.** Mean fluorescence intensity (MFI) of Ig kappa light chain (Igk) is indicative of the level of the B-cell receptor (BCR) expression at the B-cell surface. IgK MFI appears comparable for CD5$^+$CD19$^+$ tumor cells of both groups of CLL (LSR$^{Low}$, N= 4 to 6; LSR$^{High}$, N=6) and decreased compared to CD5$^-$CD19$^+$ normal B-cells. Statistical analysis was performed using Unpaired T test ns: no significant difference.
